# LotuS2: An ultrafast and highly accurate tool for amplicon sequencing analysis

**DOI:** 10.1101/2021.12.24.474111

**Authors:** Ezgi Özkurt, Joachim Fritscher, Nicola Soranzo, Duncan Y. K. Ng, Robert P. Davey, Mohammad Bahram, Falk Hildebrand

## Abstract

**Background:** Amplicon sequencing is an established and cost-efficient method for profiling microbiomes. However, many available tools to process this data require both bioinformatics skills and high computational power to process big datasets. Furthermore, there are only few tools that allow for long read amplicon data analysis. To bridge this gap, we developed the LotuS2 (Less OTU Scripts 2) pipeline, enabling user-friendly, resource friendly, and versatile analysis of raw amplicon sequences.

**Results:** In LotuS2, six different sequence clustering algorithms as well as extensive pre- and post-processing options allow for flexible data analysis by both experts, where parameters can be fully adjusted, and novices, where defaults are provided for different scenarios. We benchmarked three independent gut and soil datasets, where LotuS2 was on average 29 times faster compared to other pipelines - yet could better reproduce the alpha- and beta-diversity of technical replicate samples. Further benchmarking a mock community with known taxa composition showed that, compared to the other pipelines, LotuS2 recovered a higher fraction of correctly identified genera and species (98% and 57%, respectively). At ASV/OTU level, precision and F-score were highest for LotuS2, as was the fraction of correctly reconstructed 16S sequences.

**Conclusion:** LotuS2 is a lightweight and user-friendly pipeline that is fast, precise and streamlined. High data usage rates and reliability enable high-throughput microbiome analysis in minutes.

**Availability:** LotuS2 is available from GitHub, conda or via a Galaxy web interface, documented at http://lotus2.earlham.ac.uk/.

## BACKGROUND

The field of microbiome research has been revolutionized in the last decade, owing to methodological advances in DNA-based microbial identification. Amplicon sequencing (also known as metabarcoding) is one of the most commonly used techniques to profile microbial communities based on targeting and amplifying phylogenetically conserved genomic regions such as the 16S/18S ribosomal RNA (rRNA) or internal transcribed spacers (ITS) for identification of bacteria and eukaryotes (esp. Fungi), respectively [1,2]. The popularity of amplicon sequencing has been growing due to its broad applicability, ease-of-use, cost-efficiency, streamlined analysis workflows as well as specialist applications such as low biomass sampling [3].

Alas, amplicon sequencing comes with several technical challenges. These include primer biases [4], chimeras occurring in PCR amplifications [5], rDNA copy number variations [6] and sequencing errors that frequently inflate observed diversity [7]. Although modern read error corrections can significantly decrease artifacts of sequencing errors [8], the taxonomic resolution is limited to the genus or at best to species level [9,10]. To process amplicon sequencing data from raw reads to taxa abundance tables, several pipelines have been developed, such as mothur [11], QIIME 2 [12], DADA2 [8] or LotuS [13]. These pipelines differ in their data processing and sequence clustering strategies, reflected in differing execution speed and resulting amplicon interpretations [13,14].

Here we introduce Lotus2, designed to improve reproducibility, accuracy and ease of amplicon sequencing analysis. LotuS2 offers a completely refactored installation, including a web interface that is freely deployable on Galaxy clusters. During development, we focused on all steps of amplicon data analysis, including processing raw reads to abundance tables as well as improving taxonomic assignments and phylogenies of Operational Taxonomic Units (OTUs) or Amplicon Sequencing Variants (ASVs) at the highest quality with the latest strategies available. Pre- and post-processing steps were further improved compared to the predecessor “LotuS1”: the read filtering program sdm (simple demultiplexer) and taxonomy calculation program LCA (least common ancestor) were refactored and parallelized in C++. LotuS2 uses a ‘seed extension’ algorithm that improves the quality and length of OTU/ASV representative DNA sequences. We integrated numerous features such as additional sequence clustering options (DADA2, UNOISE3, VSEARCH and CD-HIT), advanced read quality filters based on probabilistic and Poison binomial filtering and curated ASVs/OTUs diversity and abundances (LULU, UNCROSS2, ITSx, host DNA filters). LotuS2 can also be integrated in complete workflows, e.g. the microbiome visualization-centric pipeline CoMA [15] uses LotuS1/2 at its core to estimate taxa abundances.

Here, we evaluated LotuS2 in reproducing microbiota profiles in comparison to contemporary amplicon sequencing pipelines. We found that LotuS2 consistently reproduces microbiota profiles more accurately, using three independent datasets, and reconstructs a mock community with the highest overall precision.

## MATERIALS AND METHODS

### Design Philosophy of LotuS2

Overestimating observed diversity is one of the central problems in amplicon sequencing, mainly due to sequencing errors [7,16]. The second read pair from Illumina paired-end sequencing is generally lower in quality [17] and can contain more errors than predicted from Phred quality scores alone [18,19]. Additionally, merging reads can introduce chimeras due to read pair mismatches [20]. The accumulation of errors over millions of read pairs can impact observed biodiversity, so essentially is a multiple testing problem. To avoid overestimating biodiversity, LotuS2 uses a relatively strict read filtering during the error-sensitive sequence clustering step. This is based on i) 21 quality filtering metrics (average quality, homonucleotide repeats, removal of reads without amplicon primers, etc), ii) probabilistic and Poisson binomial read filtering [17,21], iii) filtering reads that cannot be dereplicated (clustered at 100% nucleotide identity) either within or between samples and iv) using only the first read pair from paired-end Illumina sequencing platforms. These reads are termed “high-quality” reads in the pipeline description and are clustered into OTUs/ASVs, using one of the sequence clustering programs **(Figure 1B)**.

**Figure 1-.**
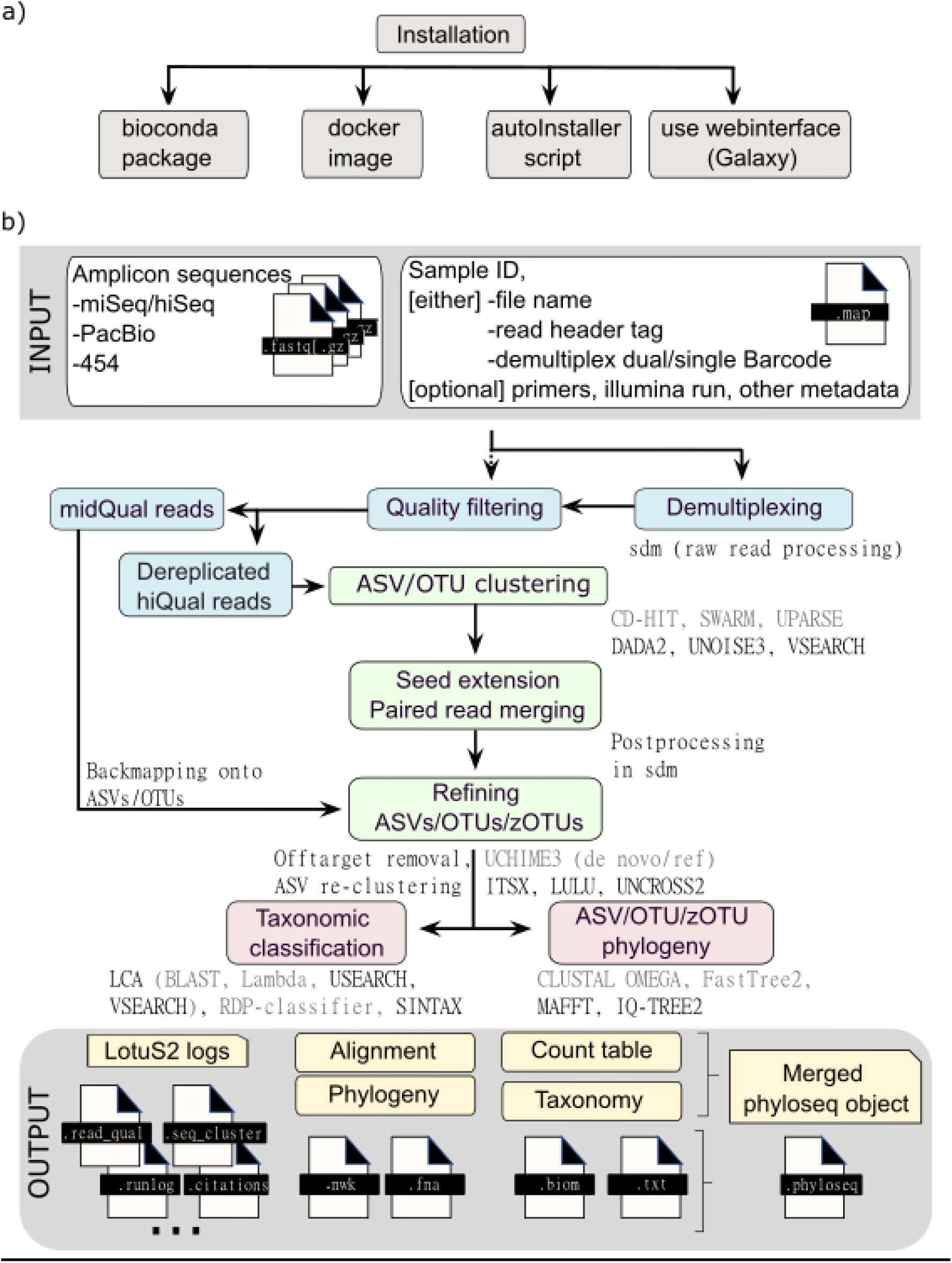
Workflow of the LotuS2 Pipeline. **a)** LotuS2 can be installed either through i) Bioconda, ii) GitHub with the provided autoInstaller script or iii) using a Docker image. Alternatively, iv) Galaxy web servers can also run LotuS2 (e.g. https://usegalaxy.eu/) **b)** LotuS2 accepts amplicon reads from different sequencing platforms, along with a map file that describes barcodes, file locations, sample IDs and other information. After demultiplexing and quality filtering, high-quality reads are clustered into either ASVs or OTUs. The optimal sequence representing each OTU/ASV is calculated in the seed extension step, where read pairs are also merged. Mid-quality reads are subsequently mapped onto these sequence clusters, to increase cluster representation in abundance matrices. From OTU/ASV sequences, a phylogenetic tree is constructed, and each cluster is taxonomically assigned. These results are made available in multiple standard formats, such as tab-delimited files, .biom or phyloseq objects, to enable downstream analysis. New options in LotuS2 for each step are denoted with black colour whereas options in grey font were already available in LotuS.

However, filtered out “mid-quality” sequences are partly recovered later in the pipeline, during the seed extension step. LotuS2 will reintroduce reads failing dereplication thresholds or being of “mid-quality” by mapping these reads back onto high-quality OTUs/ASVs if matching at ≥ 97% sequence identity. In the “seed extension” step, the optimal sequence representing each OTU/ASV is determined by comparing all (raw) reads clustered into each OTU/ASV. The best read (pair) is then selected based on the highest overall similarity to the consensus OTU/ASV, quality and length that, in the case of paired read data, can then be merged. Thereby, the seed extension step enables more reads to be included in taxa abundance estimates, as well as enabling longer ASV/OTU representative sequences to be used during taxonomic classifications and the reconstruction of a phylogenetic tree.

### Implementation of LotuS2

#### Installation

LotuS2 can be accessed either through major software repositories such as i) Bioconda, ii) as a Docker image or iii) GitHub (accessible through http://lotus2.earlham.ac.uk/) **(Figure 1A)**. The GitHub version comes with an installer script that downloads the required databases and installs and configures LotuS2 with its dependencies. Alternatively, we provide iv) a wrapper for Galaxy [22] allowing installation of LotuS2 on any Galaxy server from the Galaxy ToolShed. LotuS2 is already available to use for free on the UseGalaxy.eu server (https://usegalaxy.eu/), where raw reads can be uploaded and analysed **(Supp. Figure 1)**. While LotuS2 is natively programmed for Unix (Linux, macOS) systems, other operating systems are supported through the Docker image or the Galaxy web interface.

#### Input

LotuS2 is designed to run with a single command, where the only essential flags are the path to input files (fastq(.gz), fna(.gz) format), output directory and mapping file. The mapping file contains information on sample identifiers, demultiplexing barcodes or file paths to already demultiplexed files and can be either automatically generated or provided by the user. The sequence input is flexible, allowing simultaneous demultiplexing of read files and/or integration of already demultiplexed reads.

LotuS2 is highly configurable, enabling user-specific needs beyond the well-defined defaults. There are 63 flags that can be user-modified, including dereplication filtering thresholds (-derepMin), sequencing platform (-p), amplicon region (-amplicon_type), or OTU/ASV postprocessing (e.g. -LULU option to remove erroneous OTUs/ASVs [23]). In addition, read filtering criteria can be controlled in 32 detailed options via custom config files (defaults are provided for Illumina MiSeq, hiSeq, novaSeq, Roche 454, PacBio HiFi).

#### Output

The primary output is a set of tab-delimited OTU/ASV count tables, the phylogeny of OTUs/ASVs, their taxonomic assignments and corresponding abundance tables at different taxonomic levels. These are summarized in .biom [24] and phyloseq objects [25], that can be loaded directly by other software for downstream analysis.

Furthermore, a detailed report of each processing step can be found in the log files which contain commands of all used programs (including citations and versions) with relevant statistics. We support and encourage users to conduct further analysis in statistical programming languages such as R, Python or Matlab and using analysis packages such as phyloseq [25], documented in tutorials at http://lotus2.earlham.ac.uk/..

#### Pipeline workflow

Most of LotuS2 is implemented in PERL 5.1; computational or memory intensive components like simple demultiplexer (sdm) and LCA (least common ancestor) are implemented in C++ (see **Figure 1B** for pipeline workflow). Demultiplexing, quality filtering and dereplication of reads is implemented in sdm. Taxonomic postprocessing is implemented in LCA. Six sequence clustering methods are available: UPARSE [17], UNOISE3 [26], CD-HIT [27], SWARM [28], DADA2 [8] or VSEARCH [29].

In the “seed extension” step, a unique representative read of a sequence cluster is chosen, based on quality and merging statistics. Each sequence cluster, termed ASVs in the case of DADA2, OTUs otherwise^1^, is represented by a high confidence DNA sequence (see Design Philosophy of LotuS2 for more information).

OTUs/ASVs are further postprocessed to remove chimeras, either *de novo* and/or reference based using the program UCHIME3 [30] or VSEARCH-UCHIME [29]. By default, ITS sequences are extracted using ITSx [31]. Highly resolved OTUs/ASVs are then curated based on sequence similarity and co-occurrence patterns, using LULU [23]. False-positive OTU/ASV counts can be filtered using the UNCROSS2 algorithm [32]. OTUs/ASVs are by default aligned against the phiX genome, a synthetic genome often included in Illumina sequencing runs, using Minimap2 [33]; these OTUs/ASVs are subsequently removed. Additionally, the user can filter for host contamination by providing custom genomes (e.g., human reference), as host genome reads are often misclassified as bacterial 16S by existing pipelines [3].

Each OTU/ASV is taxonomically classified, using either RDP classifier [34], SINTAX [35] or by alignments to reference database(s), using the custom “LCA” (least common ancestor) C++ program. Alignments of OTUs/ASVs with either Lambda [36], BLAST [37], VSEARCH [29], or USEARCH [38] are compared against a user-defined range of reference databases. These databases cover the 16S, 18S, 23S, 28S rRNA gene and ITS region, by default a Lambda alignment against the SILVA database is used [39]. Other databases bundled with LotuS2 include Greengenes [40], HITdb [41], PR2 [42], beetax (bee gut-specific taxonomic annotation) [43], UNITE (fungal ITS database) [44], or users can provide reference databases (a fasta file and a tab-delimited taxonomy file). These databases can be used by themselves, or in conjunction. From mappings against one or several reference databases, the least common ancestor for each OTU/ASV is calculated using LCA. Priority is given to deeply resolved taxonomies, sorted by the earlier listed reference databases. For reconstructing phylogenetic trees, multiple sequence alignments for all OTUs/ASVs are calculated with either MAFFT [45] or Clustal Ω [46]; from these a maximum likelihood phylogeny is constructed using either fasttree2 [47] or IQ-TREE 2 [48].

### Benchmarking amplicon sequencing pipelines

To benchmark the computational performance and reproducibility, we compared LotuS2’s performance to commonly used amplicon sequencing pipelines including mothur [11], DADA2 [8], and QIIME 2 [12]. We relied, where possible, on default options or standard operating procedure (SOPs) provided by the respective developers (mothur: https://mothur.org/wiki/miseq_sop/; QIIME 2: https://docs.qiime2.org/2021.11/tutorials/moving-pictures/ and DADA2: https://benjjneb.github.io/dada2/tutorial.html). DADA2 cannot demultiplex raw reads and in these cases, LotuS2 demultiplexed raw reads were used as DADA2 input.

Our benchmarking scripts are available at https://github.com/ozkurt/lotus2_benchmarking (see **Supp. Text**). Several sequence cluster algorithms were benchmarked, for LotuS2: DADA2 [8], UPARSE [17], UNOISE3 [26], CD-HIT [27] and VSEARCH [29]; for QIIME 2: DADA2 and Deblur [49]; DADA2 supporting natively only DADA2 clustering; for mothur: OptiClust; and for LotuS1: UPARSE. For taxonomic classification, SILVA138.1 [39], was used in all pipelines. ITS amplicons are clustered with CD-HIT, UPARSE and VSEARCH and filtered by default using ITSx [31] in LotuS2. ITSx identifies likely ITS1, 5.8S and ITS2 and full-length ITS sequences, and sequences not within the confidence interval are discarded in LotuS2. In analogy, QIIME 2-DADA2 uses q2-ITSxpress [50] that also removes unlikely ITS sequences.

Error profiles during ASV clustering were inferred separately for the samples sequenced in different MiSeq runs during DADA2 and Deblur clustering in all pipelines. We truncated the reads into the same length (200 bases, default by LotuS2) in all pipelines while analysing the datasets. Primers were removed from the reads, where supported by a pipeline.

### Measuring computational performance of amplicon sequencing pipelines

When benchmarking pipelines, processing steps were separated into 5 categories in each tested pipeline: a) Pre-processing (demultiplexing if required, read filtering, primer removal and read merging for QIIME 2-Deblur), b) sequence clustering (clustering + refining of the clusters and denoising for QIIME 2-DADA2, c) OTU/ASV taxonomic assignment, d) construction of a phylogenetic tree (the option is available only in mothur, QIIME 2 and LotuS2) and e) removal of host genome (the option is available only in QIIME 2 and LotuS2). In mothur, sequence clustering and taxonomic assignment times were added since these pipeline commands are entangled (https://mothur.org/wiki/miseq_sop/).

### Data used in benchmarking pipeline performance

Four datasets with different sample characteristics (e.g., compositional complexity, target gene and region, amplicon length) were analysed: i) <u>Gut-16S</u> dataset [13]: 16S rRNA gene amplicon sequencing of 40 human faecal samples in technical replicates that were sequenced in separate MiSeq runs, totalling 35,412,313 paired-end reads. Technical replicates were created by extracting DNA twice from each faecal sample. Since the Illumina runs were not demultiplexed, pipelines had to demultiplex these sequences, if available. ii) <u>Soil-16S</u> dataset: 16S rRNA gene amplicon sequencing of two technical replicates (single DNA extraction per sample) from 50 soil samples, that were sequenced in separate MiSeq runs, totalling 11,820,327 paired-end reads. PCR reactions were conducted using the 16S rRNA region primers 515F (GTGYCAGCMGCCGCGGTAA) and 926R (GGCCGYCAATTYMTTTRAGTTT). The soil-16S dataset was already demultiplexed, requiring pipelines to work with paired FASTQ files per sample. iii) <u>Soil-ITS</u> dataset: ITS amplicon sequencing of 50 technical replicates of soil samples (single DNA extraction per sample), sequenced in two independent Illumina MiSeq runs, totalling 6,006,089 paired-end reads. ITS region primers gITS7ngs_201 (GGGTGARTCATCRARTYTTTG) and ITS4ngsUni_201 (CCTSCSCTTANTDATATGC) [51] were used to amplify DNA extracted from soil samples. The soil-ITS dataset was already demultiplexed.

iv) <u>Mock</u> dataset [52]: A microbial mock community with known species composition, *mock-16* [52]. The mock dataset comprised a total of 59 strains of Bacteria and Archaea, representing 35 bacterial and 8 archaeal genera. The mock community was sequenced on an Illumina MiSeq (paired-end) by targeting the V4 region of the 16S rRNA gene using the primers 515F (GTGCCAGCMGCCGCGGTAA) and 806R (GGACTACHVGGGTWTCTAAT) [52]. This dataset was demultiplexed and contained 593,868 paired reads.

### Benchmarking the computational performance of amplicon sequencing pipelines

To evaluate the computational performance of LotuS2 in comparison to QIIME 2 [12], DADA2 [8], and the last released version of LotuS [13] (v1.62 from Jan 2020; called LotuS1 here), all pipelines were run with 12 threads on a single computer free of other workloads (CPU: Intel(R) Xeon(R) Gold 6130 CPU @ 2.10 GHz, 32 cores, 375 GB RAM). To reduce the influence of network latencies on pipeline execution, all temporary, input, and output data were stored on a local SSD. Each pipeline was run three times consecutively to account for pre-cached data and to obtain average execution time and maximum memory usage. To calculate the fold differences in execution speed between pipelines, the average time of all LotuS2 runs was divided by average QIIME 2, mothur and DADA2, where used in each of the three non-mock datasets. The average of these numbers was used to estimate the average speed advantage of LotuS2.

### Benchmarking reproducibility of amplicon sequencing pipelines

Technical replicates of the soil and gut samples were used to estimate the reproducibility of the microbial community composition between replicates. This was measured by calculating beta and alpha diversity differences between technical replicate samples. To calculate beta diversity, either Jaccard (measuring presence/absence of OTUs/ASVs) or Bray-Curtis dissimilarity (measuring both presence/absence and abundances of OTUs/ASVs) were computed between technical replicate samples. Before computing Bray-Curtis distances, abundance matrices were normalized. Jaccard distances between samples were calculated by first rarefying abundance matrices to an equal number of reads (to the size of the first sample having > 1000 read counts) per sample using RTK [53]. Significance of pairwise comparisons of the pipelines in beta diversity differences was calculated using the ANOVA test where Tukey’s HSD (honest significant differences) test was used as a *post hoc* test in R.

To calculate alpha diversity, abundance data were first rarefied to an equal number of reads per sample. Significance of each pairwise comparison in alpha diversity was calculated based on a paired Wilcoxon test, pairing technical replicates.

### Analysis of the mock community

We used an already sequenced mock community [52] of known relative composition and with sequenced reference genomes available. Firstly, taxonomic abundance tables (taxonomic assignments based on SILVA 138.1 [39] in all pipelines) were compared to the expected taxonomic composition of the sequenced mock community. Precision was calculated as (TP/(TP+FP)), recall as (TP/(TP+FN)) and F-score as (2*precision*recall/(precision+recall)), TP (true positive) being taxa present in the mock and correctly identified as present, FN (false negative) being taxa present in the mock but not identified as present and FP (false positive) being taxa absent in the mock but identified as present. The fraction of read counts assigned to true positive taxa was calculated based on the sum of the relative abundance of all true positive taxa. These scores were calculated at different taxonomic levels.

Secondly, we investigated the precision of reconstructed 16S rRNA nucleotide sequences, representing each OTU or ASV, by calculating the nucleotide similarity between ASVs/OTUs and the known reference 16S rRNA sequences. To obtain the nucleotide similarity, we aligned ASV/OTU DNA sequences from tested pipelines via BLAST to a custom reference database that contained the 16S rRNA gene sequences from the mock community (https://github.com/caporaso-lab/mockrobiota/blob/master/data/mock-16/source/expected-sequences.fasta), using the –taxOnly option from LotuS2. The BLAST % nucleotide identity was subsequently used to calculate the best matching 16S rRNA sequence per ASV/OTU.

## RESULTS

We analysed four datasets to benchmark the computational performance and reliability of the pipelines. The datasets consisted either of technical replicates (gut-16S, soil-16S, soil-ITS) or a mock community. Technical replicates were used to evaluate the reproducibility of community structures and were chosen to represent different biomes (gut, soil), using different 16S rRNA amplicon primers (gut-16S, soil-16S) or ITS sequences (soil-ITS) as well as a synthetic mock community of known composition.

### Computational performance and data usage

The complete analysis of the gut-16S dataset was fastest in LotuS2 (on average 35, 12, 9 and 3.8 times faster than mothur, QIIME 2-DADA2, QIIME 2-DEBLUR and native DADA2, respectively, **Figure 2A)**. Note that DADA2 could not demultiplex the dataset, the average of LotuS2 and QIIME2 demultiplexing times were used instead. LotuS2 was also faster in the analysis of the soil-16S dataset compared to the other tested pipelines (5.7, 3.5, 3.5 times faster than DADA2, QIIME 2-DADA2 and QIIME 2-DEBLUR, respectively, **Figure 2B**). The difference in speed between LotuS2 and QIIME 2 was more pronounced in the analysis of the soil-ITS dataset, where LotuS2 was on average 69 times faster than QIIME 2 and DADA2 **(Figure 2C)**. LotuS2 also outperformed other pipelines in the case of the gut-16S dataset (on average LotuS2 was 15 times faster) compared to the soil dataset (average 4.2). This difference stems mainly from the demultiplexing step, where LotuS2 is significantly faster. The sequence clustering step was fastest using the UPARSE algorithm, i.e. an average 60-fold faster than sequence clustering in other pipelines. Averaged over these three datasets, LotuS2 was 29 times faster than other pipelines.

**Figure 2:**
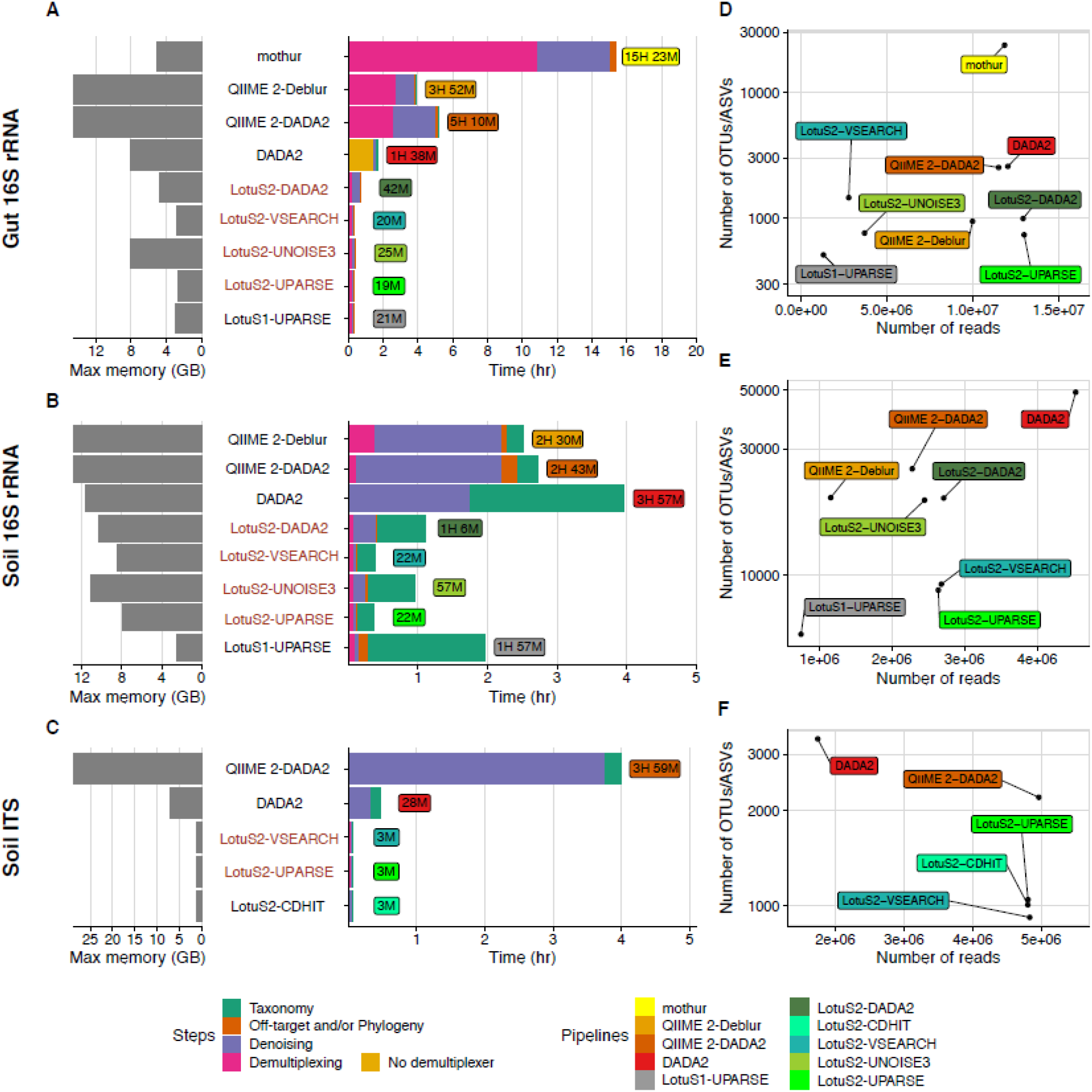
Computational performance of amplicon sequencing pipelines. 16S rRNA amplicon MiSeq data from A) gut-16S and B) soil-16S and C) soil-ITS samples were processed to benchmark resource usage of each pipeline, run on the same system under equal conditions (12 cores, max 150Gb memory). In all pipelines, OTUs/ASVs were classified by similarity comparisons to SILVA 138.1. In LotuS2, LAMBDA was used to align sequences for all clustering algorithms. Pipeline runs were separated by common steps (pre-processing, sequence clustering, taxonomic classification and phylogenetic tree construction and/or off-target removal). Because native DADA2 cannot demultiplex reads, we used the average demultiplexing time of QIIME 2 and LotuS2 (LotuS2 demultiplexed, unfiltered reads were provided to DADA2). LotuS2 pipelines are labelled with red colour. D, E, F) Data usage efficiency of each tested pipeline, by comparing the number of sequence clusters (OTUs or ASVs) to retrieved read counts in the final output matrix of each pipeline. Note that mothur results on soil-16S are not shown, because the pipeline rejected with default parameters all sequences.

Taxonomic classification of OTUs/ASVs was also faster in LotuS2 (~5 times faster for gut-16S, 2 times for soil-16S). However, this strongly depends on the total number of OTUs/ASVs for all pipelines. For example, the default naïve-Bayes classifier [54] in QIIME 2 is faster relative to the number of OTUs/ASVs, compared to LotuS2 taxonomic assignments in this benchmark. Nevertheless, the LotuS2 default taxonomic classification is via RDP classifier [34], and alternatively, the SINTAX [35] classifier could be used, both of which are significantly faster than the here presented Lambda LCA against the Silva reference database.

Compared to LotuS1, LotuS2 was on average 3.2 times faster, likely related to refactored C++ programs that can take advantage of multiple CPU threads **(Figure 2A-B)**.

In its fastest configuration (using “UPARSE” option in clustering, “RDP” to assign taxonomy), the gut and soil 16S rRNA datasets can be processed with LotuS2 in under 20 mins and 12 mins, using < 10 GB of memory and 4 CPU cores.

Despite using similar clustering algorithms (e.g. DADA2 is used in DADA2, QIIME 2 and LotuS2), the tested pipelines apply different pre- and post-processing algorithms to raw sequence reads and clustered ASVs and OTUs, leading to differing ASV/OTU numbers and retrieved reads (the total read count in the ASV/OTU abundance matrix) **(Supp. Table 1 and Figure 2D-F)**. DADA2 typically estimated the highest number of ASVs, but the number of retrieved reads varied strongly between datasets. QIIME 2-DADA2 estimated fewer ASVs than DADA2, but more ASVs than LotuS2-DADA2, although mapping fewer reads than LotuS2. Although retrieving a smaller number of reads, QIIME 2-Deblur reported comparable numbers of ASVs to LotuS2, despite the differences in clustering algorithms. mothur performed differently in the gut-16S and soil-16S datasets, where it estimated either the highest number of OTUs or could not complete the analysis since all the reads being filtered out, respectively. Overall, LotuS2 often reported the fewest ASVs/OTUs, while including more sequence reads in abundance tables. This indicates that LotuS2 has a more efficient usage of input data while covering a larger sequence space per ASV/OTU.

### Benchmarking the reproducibility of community compositions

Next, we assessed the reproducibility of community compositions, using gut-16S, soil-16S and soil-ITS datasets comparing beta diversity between technical replicates (Bray Curtis distance, BCd and Jaccard distance, Jd). We found that Jd and BCd were the lowest in LotuS2, largely independent of the chosen sequence clustering algorithms and dataset. This indicates a greater reproducibility of community compositions generated by LotuS2 **(Figure 3A-B and Supp. Figure 2**). The lowest BCd and Jd were observed for UPARSE **(Figure 3A-B and Supp. Figure 2)** in both gut- and soil-16S datasets, though this was not always significant between different LotuS2 runs **(Supp. Table 2)**.

**Figure 3-.**
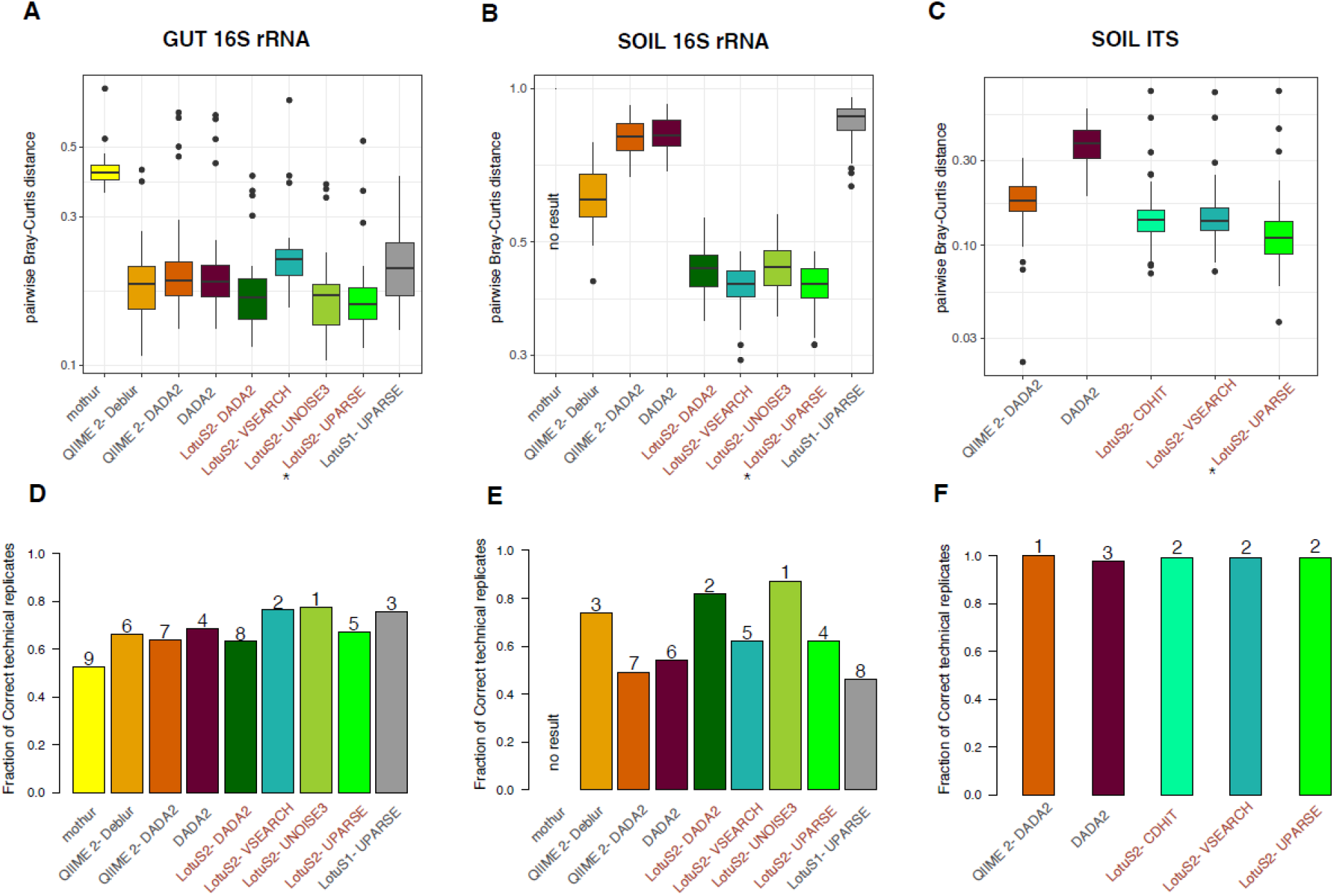
Reproducibility from different amplicons sequence data analysis pipelines. Three independent datasets were used to represent different biomes and amplicon technologies, using A, D) human faecal samples (16S rRNA gene, N=40 replicates). B, E) soil samples (16S rRNA gene, N=50 replicates) and C, F) soil samples (ITS 2, N=50 replicates). A-C) Bray-Curtis distances among technical replicate samples are used to assess the reproducibility of community compositions by different pipelines. The pipeline with the lowest BCd in each subfigure is denoted with a star (*). The significance of pairwise comparisons of each pipeline is calculated using the Tukey’s HSD test **(Supp. Table 2)**. D-F) Further, the fraction of technical replicates being closest to each other (BCd) was calculated to simulate identifying technical replicates without additional knowledge. Numbers above bars are the ordered pipelines performing best. Lower Bray-Curtis distances between technical replicates and a higher fraction of correct technical replicates indicate better reproducibility. LotuS2 pipelines are labelled with red colour.

Even using the same clustering algorithm, LotuS2-DADA2 compositions were more reproducible, compared to both QIIME 2-DADA2 and DADA2 (significant only on soil data). LotuS2-DADA2 denoises by default all reads (per sequencing run) together, while in the default DADA2 setup each sample is separately denoised; the latter strategy has a reduced computational burden but can potentially miss sequence information from rare bacteria. mothur showed poorer performance compared to other pipelines on the gut-16S dataset and did not complete on the soil data.

We then calculated the fraction of samples being closest in BCd distance to its technical replicate for each pipeline **(Figure 3D-E)**, simulating the process of identifying technical replicates without prior knowledge. LotuS2 with UNOISE3 clustering resulted in the highest fraction of samples being closest to its replicate among all samples, in both gut- and soil-16S datasets while in the mothur result, technical replicates were the most unlikely to be closest to their technical replicate.

When this comparison was made with the non-default options in LotuS2 (using different dereplication parameters, deactivating LULU, using UNCROSS2 or retaining taxonomically unclassified reads), BCd between the technical replicates remained largely unchanged **(Supp. Figure 2, Supp. Figure 3A-B and Supp. Text)**. However, retaining unclassified reads could significantly reduce the reproducibility of LotuS2 results on the gut-16S dataset. Furthermore, even starting the analysis with different read truncation lengths, LotuS2 still had the highest reproducibility in both gut- and soil-16S datasets **(Supp. Figure 4, Supp. Figure 5 and Supp. Text)**.

Lastly, we calculated the reproducibility of reported alpha diversity between technical replicate samples in both gut-16S and soil-16S datasets **(Supp. Figure 6A-B)**. In both datasets, LotuS2 alpha diversity was not significantly different between technical replicates, as expected (5 of 8 comparisons, Wilcoxon signed-rank test), whereas, in 6 of 6 cases, QIIME 2, mothur and DADA2 had significant differences in the alpha diversity between technical replicates. Thus, LotuS2 showed in our benchmarks a higher Thus, LotuS2 showed in our benchmarks a higher data usage efficiency and higher reproducibility of community compositions than QIIME 2, DADA2 and mothur. These benchmarks also showed the importance of pre- and postprocessing raw reads and OTUs/ASVs, since LotuS2-DADA2 and QIIME 2-DADA2 performed better than and DADA2, despite using the same clustering algorithm.

### Benchmarking soil-ITS dataset

Unlike 16S rRNA gene amplicons, ITS amplicons typically vary greatly in length [4], requiring a different sequence clustering workflow; therefore, LotuS2 uses by default CD-HIT to cluster ITS sequences, and ITSx to identify plausible ITS1/2 sequences.

In terms of data usage, both LotuS2 and QIIME 2-DADA2 retrieved similar numbers of reads, but for QIIME 2 these read counts were distributed across twice the number of ASVs **(Figure 2F)**. QIIME 2-DADA2 reproduced the fungal composition significantly worse in replicate samples, compared to LotuS2-UPARSE, having higher pairwise BCd **(Figure 3C)** and Jd **(Supp. Figure 2H-I)**. However, it spanned the highest fraction of samples closest to its technical replicate, although this fraction was overall very high for all the pipelines (0.978-1) **(Figure 3F)**. DADA2 performed relatively worse, yielding the highest number of ASV, lowest retrieved read counts **(Figure 2F)**, significantly the highest BCd **(Figure 3C, Suppl. Table 2)** between replicate samples. LotuS2 had overall the lowest BCd and Jd between replicates, using both UPARSE and CD-HIT clustering **(Figure 3C, Supp. Figure 2H-I)**. Usage of CD-HIT in combination with ITSx led to an increase in OTU diversity (from 947 to 1008) although read counts remained mostly the same in the final output matrix and BCd was largely similar **(Supp. Figure 3C)**. Here, deactivating LULU slightly decreased reproducibility **(Supp. Figure 3C)**.

Finally, we calculated the reproducibility of alpha diversity between the technical replicate samples in the soil-ITS dataset **(Supp. Figure 6C)**. All pipelines resulted in no significant difference between the technical replicate samples, thus alpha diversity was highly reproducible independent of the pipeline.

### Benchmarking the dataset from the mock microbial community

To assess how well a known community can be reconstructed in LotuS2, we used a previously sequenced 16S mock community [52] containing 43 genera and 59 microbial strains, where complete reference genomes were available.

All pipelines performed poorly at reconstructing the community composition (Pearson R=0.43-0.67, Spearman Rho=0.54-0.80, **Supp. Table 3 and Supp. Figure 7**), possibly related to PCR biases and rRNA gene copy number variation. Therefore, we focused on the number of correctly identified taxa. For this, we calculated the number of reads assigned to true taxa as well as precision, recall and F-score at genus level. LotuS2-VSEARCH and LotuS2-UPARSE had the highest precision, F-score and fraction of reads assigned true positive taxa, **(Figure 4A and Supp. Figure 8)**. LotuS1 had the highest recall, but low precision. When applying the same tests at species level, LotuS2-DADA2 had overall the highest precision and F-score **(Supp. Figure 9)**. QIIME 2-DEBLUR had often competitive, but slightly lower, precision, recall and F-scores compared to LotuS2, while mothur, QIIME 2-DADA2 and DADA2 scores were lower **(Figure 4A)**.

**Figure 4-.**
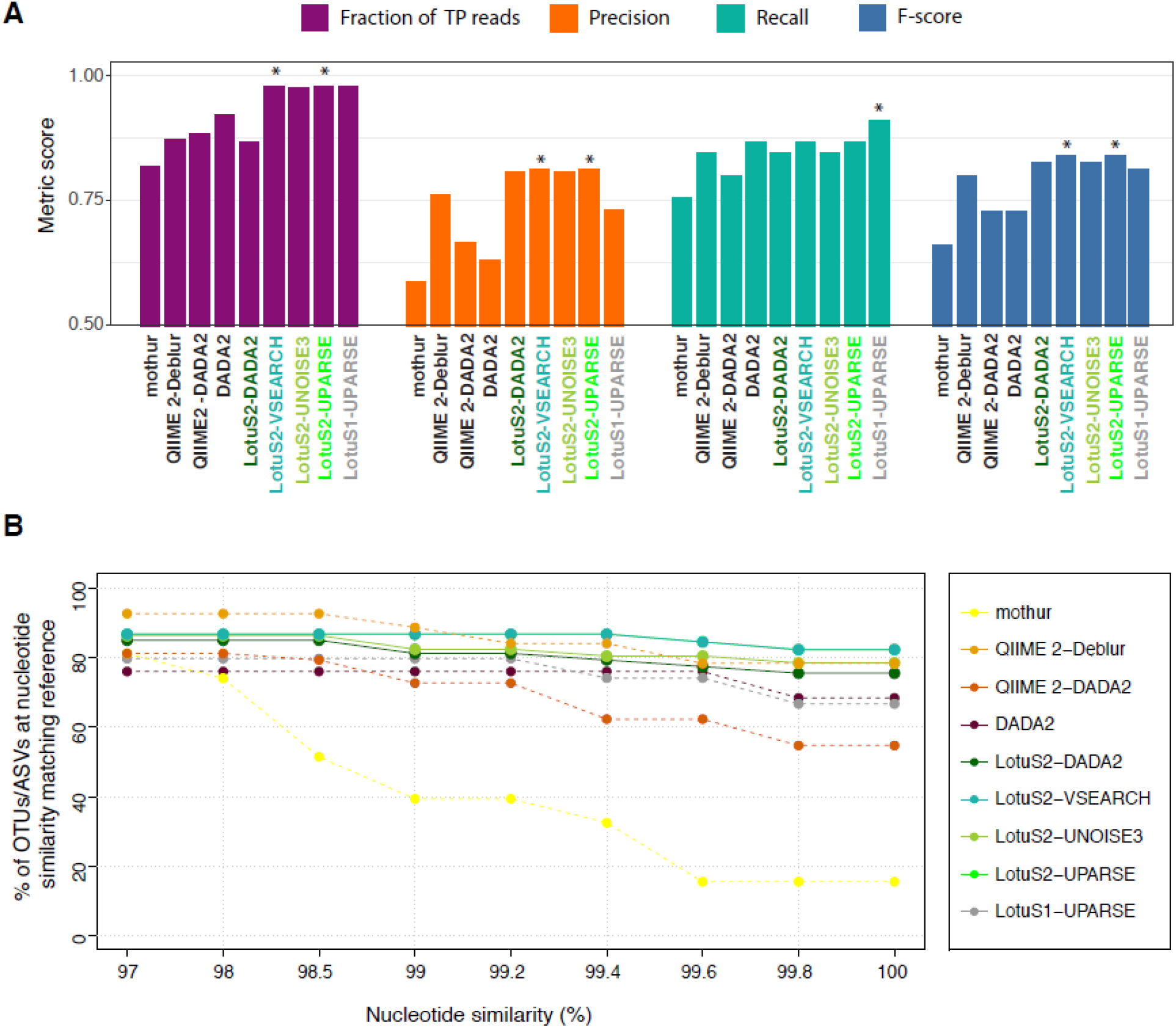
Benchmarking of amplicon sequence data analysis pipeline’s performance using a mock community with known species composition. A) Accuracy of each pipeline in predicting the mock community composition at genus level. For benchmarking we compared the fraction of reads assigned to true genera and both correctly and erroneously recovered genera. Precision, Recall and F-score were calculated based on the true positive, false positive and false negative taxa identified. At species level, LotuS2 excelled as well in these statistics (Supp. Figure 9). B) Percentage of true positive ASVs/OTUs having a nucleotide identity ≥ indicated thresholds to 16S rRNA gene sequences of genomes from the mock community. Pipeline(s) showing the highest performance in each comparison is denoted with a star (*). TP, true positive; ASV, amplicon sequencing variant; OTU, operational taxonomic unit.

Next, we investigated which software could best reconstruct the correct OTU/ASV sequences. For this, we calculated the fraction of TP OTUs/ASVs (i.e., OTUs/ASVs which are assigned to a species based on the custom mock reference taxonomy) with 97%-100% nucleotide identity to 16S rRNA sequences from reference genomes in each pipeline **(Figure 4B)**. Here, LotuS2-VSEARCH and LotuS2-UPARSE reconstructed OTU sequences were most often identical to the expected sequences, having 82.2% of the OTU sequences reconstructed at 100% nucleotide identity to reference sequences. QIIME 2-Deblur ASV sequences were of similar quality, but slightly less often at 100% nucleotide identity (78.2%). DADA2 and QIIME 2-DADA2 ASV sequences were often more dissimilar to the expected reference sequences. It is noteworthy that LotuS2-DADA2 did outperform these two pipelines based on the same sequence clustering algorithm, likely related to the stringent read filtering and seed extension step in LotuS2.

The mock community consisted of 49 bacteria and 10 archaea [52], with 128 16S rRNA gene copies included in their genomes. If multiple 16S copies occur within a single genome, these can diverge but are mostly highly similar or even identical to each other [55]. Thus, 59 OTUs would be the expected biodiversity, and ≤128 ASVs. Notably, the number of mothur and QIIME 2-Deblur TP ASVs/OTUs exceeded this threshold (N=370, 198, respectively), both pipelines overestimate known biodiversity. DADA2 and QIIME 2-DADA2 generated more ASVs than expected per species (N=94, 122 respectively), but this might account for divergent within-genome 16S rRNA gene copies. LotuS2 was notably at the lower end in predicted biodiversity, predicting between 53-61 OTUs or ASVs in different clustering algorithms **(Supp. Table 4)**. However, these seemed to mostly represent single species, covering the present species best among pipelines, as the precision at species level was highest for LotuS2 **(Supp. Figure 9)**, thus capturing species level biodiversity most accurately.

Based on the mock community data LotuS2 was more precise in reconstructing 16S rRNA gene sequences, assigning the correct taxonomy, detecting biodiversity, and within-genome 16S copies were less likely to be clustered separately using LotuS2.

## DISCUSSION

LotuS2 offers a fast, accurate and streamlined amplicon data analysis with new features and substantial improvements since LotuS1. Software and workflow optimizations make LotuS2 substantially faster than either QIIME 2, DADA2 and mothur. On large datasets, this advantage becomes crucial for users: for example, we processed a highly diverse soil dataset consisting of >11 million non-demultiplexed PacBio HiFi amplicons (26 Sequel II libraries) in 2.5 days on 16 CPU cores, using a single command (unpublished data). Besides being more resource and user-friendly, compositional matrices from LotuS2 were more reproducible and accurate across all tested datasets (gut 16S, soil 16S, soil ITS, mock community 16S).

LotuS2 owes high reproducibility and accuracy to the efficient use of reads based on their quality tiers in different steps of the pipeline. Low-quality reads introduce noise and can artificially inflate observed biodiversity, i.e., the number of OTUs/ASVs [56]. Conversely, an overly strict read filter will decrease sensitivity for low-abundant members of a community by artificially reducing sequencing depth. To find a trade-off, LotuS2 uses only truncated, high-quality reads for sequence clustering (except ITS amplicons), while the read backmapping and seed extension steps restore some of the discarded sequence data.

Notably, OTU/ASV reconstructed with LotuS2 were the most similar (at >99% identity) to the reference, compared to other pipelines **(Figure 4B)**. This was mostly independent of clustering algorithms used, a combination of both selecting high-quality reads for sequence clustering and the seed extension step, that selects a high-quality read (pair) best representing each OTU or ASV. Seed extension also decouples read clustering and read merging, avoiding the use of the error-prone 3’ read end or second read pair during the error sensitive sequence clustering step [17]. Thereby, potential length restrictions during the clustering step will not carry over to computational steps benefitting from longer sequences, such as taxonomic assignments or phylogeny reconstructions.

In conclusion, LotuS2 is a major improvement over LotuS1, representing pipeline updates that accumulated over the past eight years. It offers superior computational performance, accuracy and reproducibility of results, compared to the other tested pipelines. Importantly, it is straightforward to install, and programmed to reduce required user time and knowledge, following the idea that less is more with LotuS2.

## Availability and Requirements

**Availability of LotuS2: Documentation, tutorials**: lotus2.earlham.ac.uk, <u>Installation via bioconda</u>: https://anaconda.org/bioconda/lotus2

Galaxy wrapper (MIT licensed): https://github.com/TGAC/earlham-galaxytools/tree/master/tools/lotus2 and https://toolshed.g2.bx.psu.edu/view/earlhaminst/lotus2/ <u>Galaxy server</u>: https://usegalaxy.eu/

Programs (GPLv3 licensed): https://github.com/hildebra/lotus2, https://github.com/hildebra/sdm, https://github.com/hildebra/LCA

All the commands used for the benchmarking are available in https://github.com/okurt/lotus2_benchmarking

## Availability of the data

Accession numbers for the datasets used for benchmarking in this study are: PRJEB49356 Mock-16 community is downloaded from the *mockrobiota* repository [52]: https://s3-us-west-2.amazonaws.com/mockrobiota/latest/mock-16/mock-forward-read.fastq.gz https://s3-us-west-2.amazonaws.com/mockrobiota/latest/mock-16/mock-reverse-read.fastq.gz

## List of abbreviations

OTU: Operational taxonomic unit
ASV: Amplicon sequence variant
ITS: Internal transcribed spacer
TP: True positive
FN: False negative
FP: False positive
LotuS: Less OTU Scripts
sdm: simple demultiplexer
LCA: least common ancestor
DADA: The **Divisive** Amplicon Denoising Algorithm
QIIME: Quantitative Insights Into Microbial Ecology

## Author contributions

FH programmed LotuS2, sdm and LCA with contributions from JF, EO, MB and NS. EO benchmarked pipelines with help from FH and DN. Websites, Galaxy interface, conda support and installation scripts for LotuS2 were implemented by FH, JF, NS and EO. EO and FH wrote the manuscript with contributions from all authors.

## Acknowledgements

The authors gratefully thank numerous LotuS1 users for consistent feedback and suggestions over the years, Sarah Worsley for her valuable comments on the manuscript and Stefano Romano and Rebecca Ansorge for their user-comments on LotuS2. We also would like to acknowledge CyVerse UK for the hosting of the LotuS2 website.

## Supp. Figures and Tables

**Supp. Figure 1:**
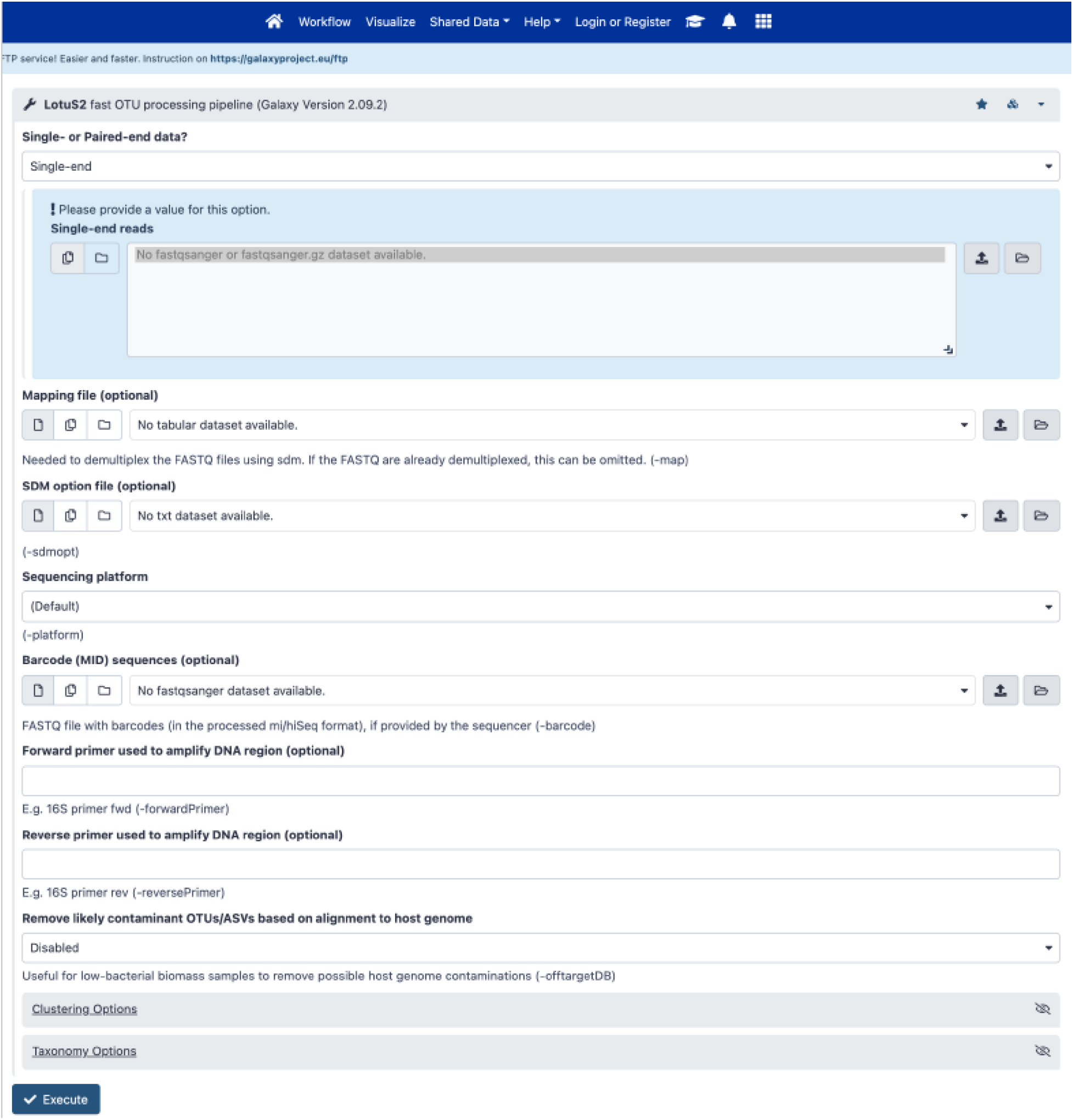
Galaxy web interface of LotuS2. Raw reads can be uploaded into the LotuS2 via the Galaxy web interface and analysed (accessible on https://usegalaxy.eu/).

**Supp. Figure 2-.**
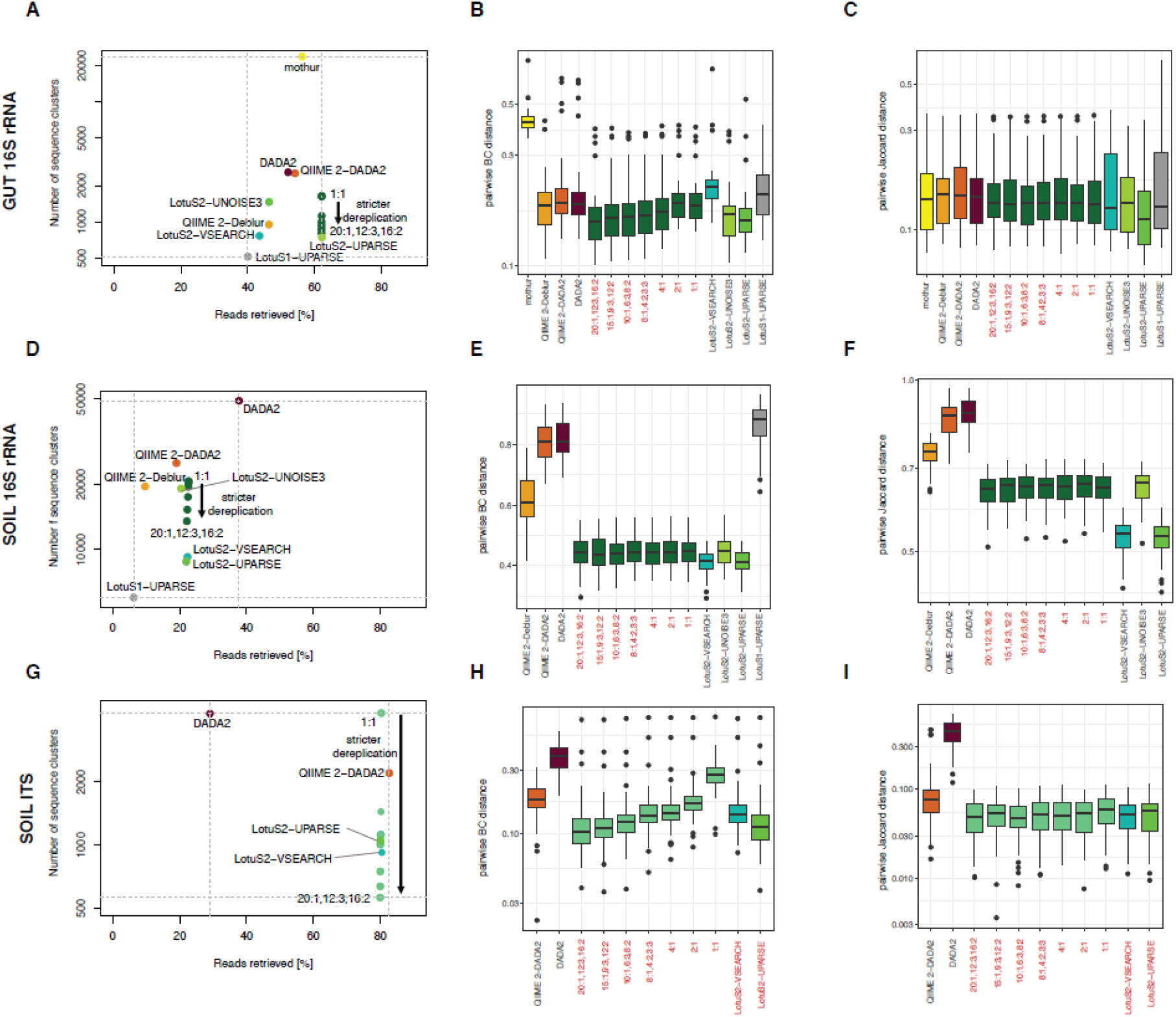
Reproducibility and data usage efficiency respective to dereplication filtering. A, D and G) Data usage efficiency of each tested pipeline at different dereplication parameters of LotuS2 (from strictest to least strict dereplication: 20:1,12:3,6:2; 15:1,9:3,12:2; 10:1,6:3,8:2; 8:1,4:2,3:3 (default); 4:1; 2:1 and 1:1) using DADA2 or CD-HIT clustering for 16S and ITS dataset, respectively, by comparing the number of sequence clusters (OTUs/ASVs) to retrieved read counts in final output matrix. The dereplication can be fine controlled through a syntax. For example, 8:1,4:2,3:3 means that a read is accepted, if it occurs >=8 times in >= 1 samples **or** >4 times total in >= 2 samples **or** >=3 times in >= 3 samples.

**Supp. Figure 3-.**
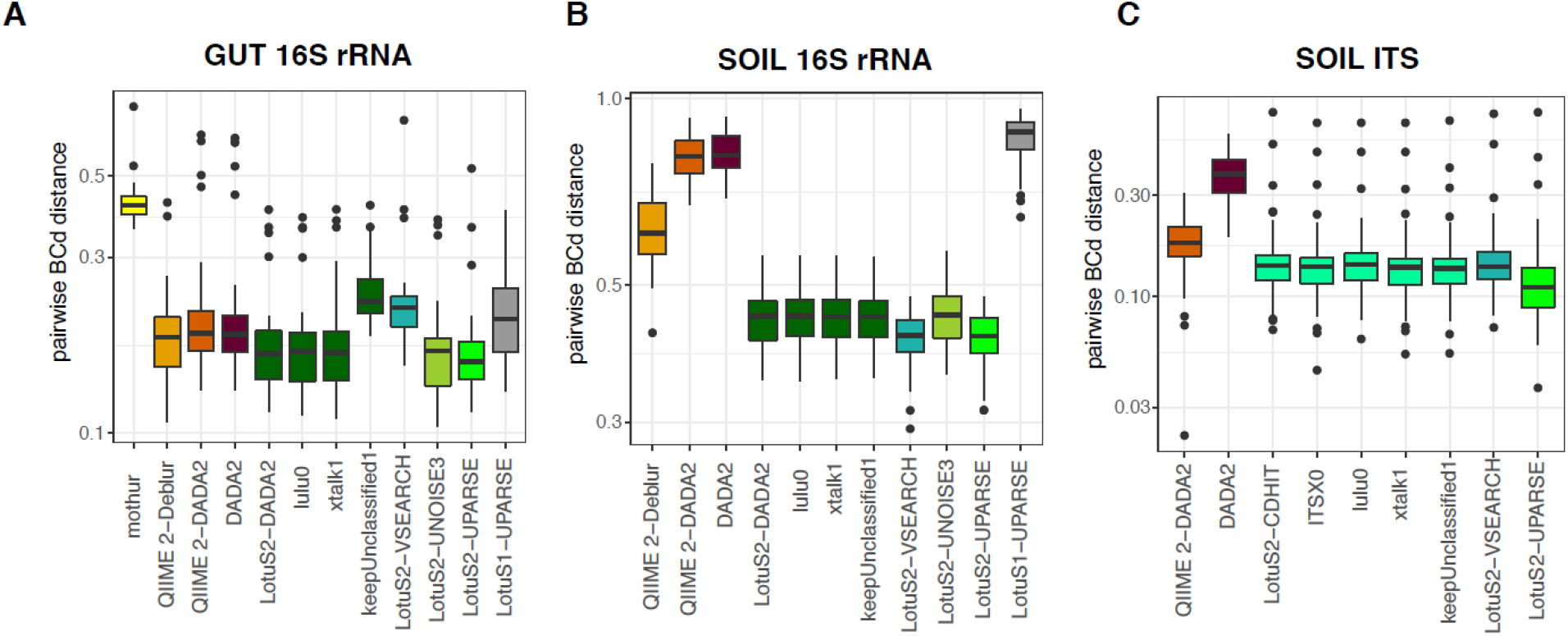
Reproducibility of the technical replicates respective to different LotuS2 non-default parameters. Bray-Curtis distances between technical replicates of A) gut-16S B) soil-16S and C) soil-ITS datasets using default and non-default parameters (LotuS2 flags: -lulu 0, -xtalk 1, -keepUnclassified 1, -ITSx 0, where 1 means the option is activated; 0 means deactivated). When activated, -lulu option uses LULU R package [23] to merge OTUs/ASVs based on their co-occurrences; -xtalk option checks for cross-talk [32], -keepUnclassified includes unclassified (i.e. not matching to any taxon in the taxonomy database) OTUs/ASVs in the final matrix and – ITSx activates the ITSx program [31] to only retain OTUs fitting to ITS1/ITS2 hmm models.

**Supp. Figure 4-.**
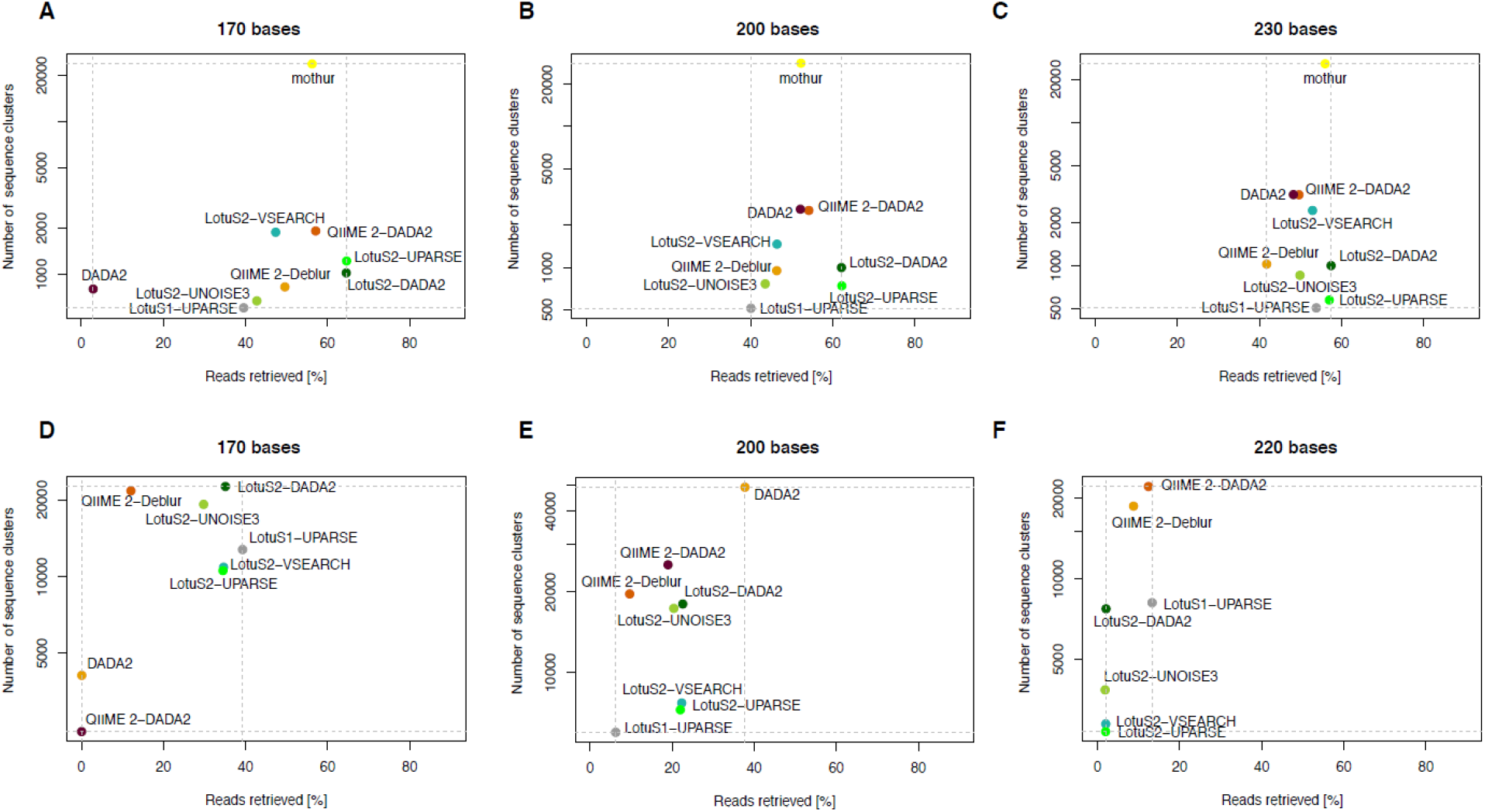
Data usage efficiency of different amplicon sequence data analysis pipelines. Data usage efficiency on gut 16S rRNA (gut-16S), soil 16S rRNA (soil-16S) and Soil ITS (soil-ITS) amplicons, tested with different pipelines at different read truncation lengths (170, 200, 230 & 170, 200, 220 bases for the gut and soil datasets, respectively), by comparing the number of sequence clusters (ASVs /OTUs) to retrieved read counts in the final output matrix of each pipeline. In all other analysis, default values were used for LotuS2 (200 bases).

**Supp. Figure 5-.**
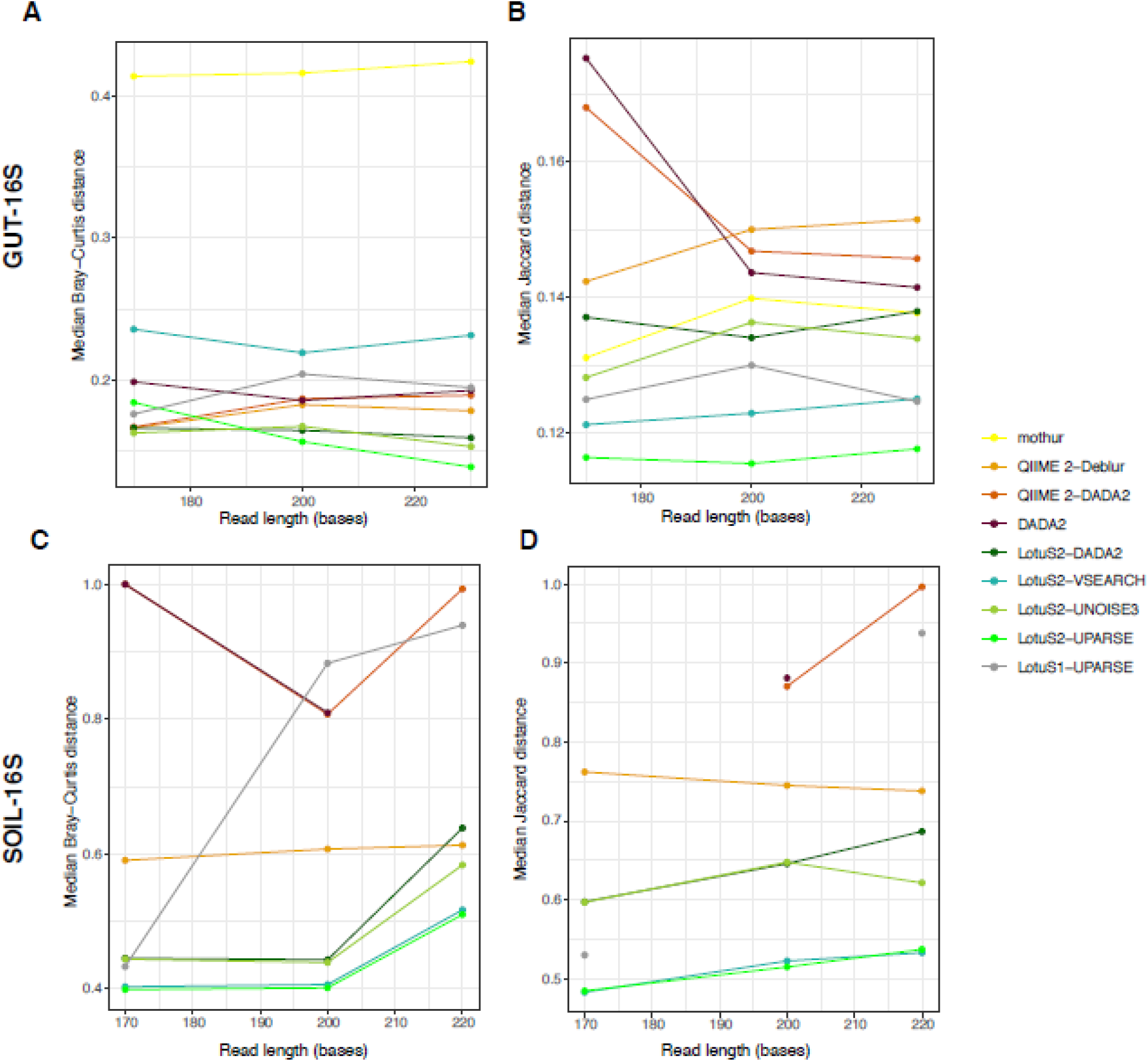
Reproducibility of beta diversity at different read truncation lengths. Reproducibility of sequenced technical replicates, by measuring the Bray-Curtis (A and C) and Jaccard distances (B and D) of the microbiome composition among technical replicate samples. Two datasets were used to represent different biomes and amplicon technologies, using (A, B) and human faecal samples (16S rRNA primer, N=40 replicates) and (C, D) soil samples (16S rRNA, V4-V5 region primers, N=50 replicates). Lower Bray-Curtis or Jaccard distances between technical replicates indicate better reproducibility of community compositions. Default pipeline parameters and recommended settings for each dataset were used (Please see the Supp. Text for further information).

**Supp. Figure 6:**
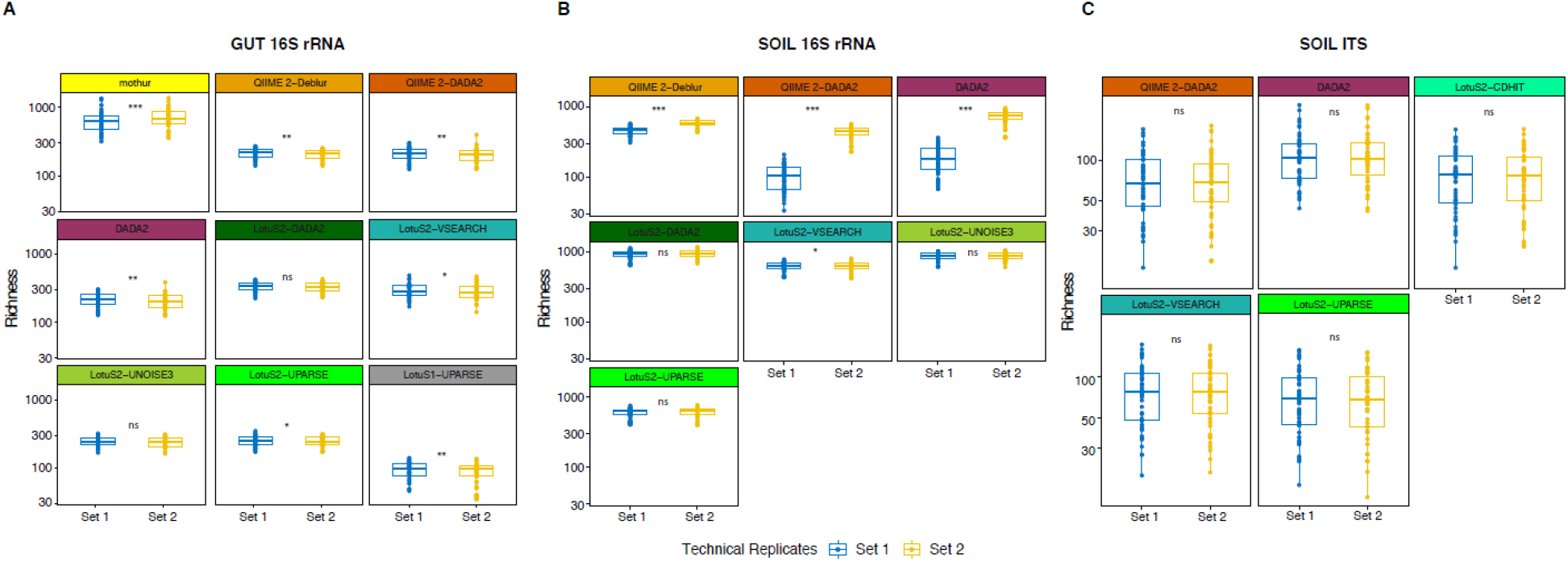
Reproducibility of alpha diversity between technical replicates. OTU/ASV Richness was calculated for A) gut-16S B) soil-16S and C) soil-ITS datasets. Samples were rarefied to an equal number of reads per sample before calculating richness, and any samples whose replicate pair was removed after rarefaction (because of having lower number of reads than the rarefaction depth) were excluded from further analysis. LotuS1 results for soil-16S were removed due to too many samples being removed in rarefactions. Significance of differences in richness between the sets were calculated based on the paired samples Wilcoxon test (***, **, * and “ns” denotes p<0.0005, p<0.005, p<0.05 and p> 0.05 (i.e. not significant), respectively).

**Supp. Figure 7:**
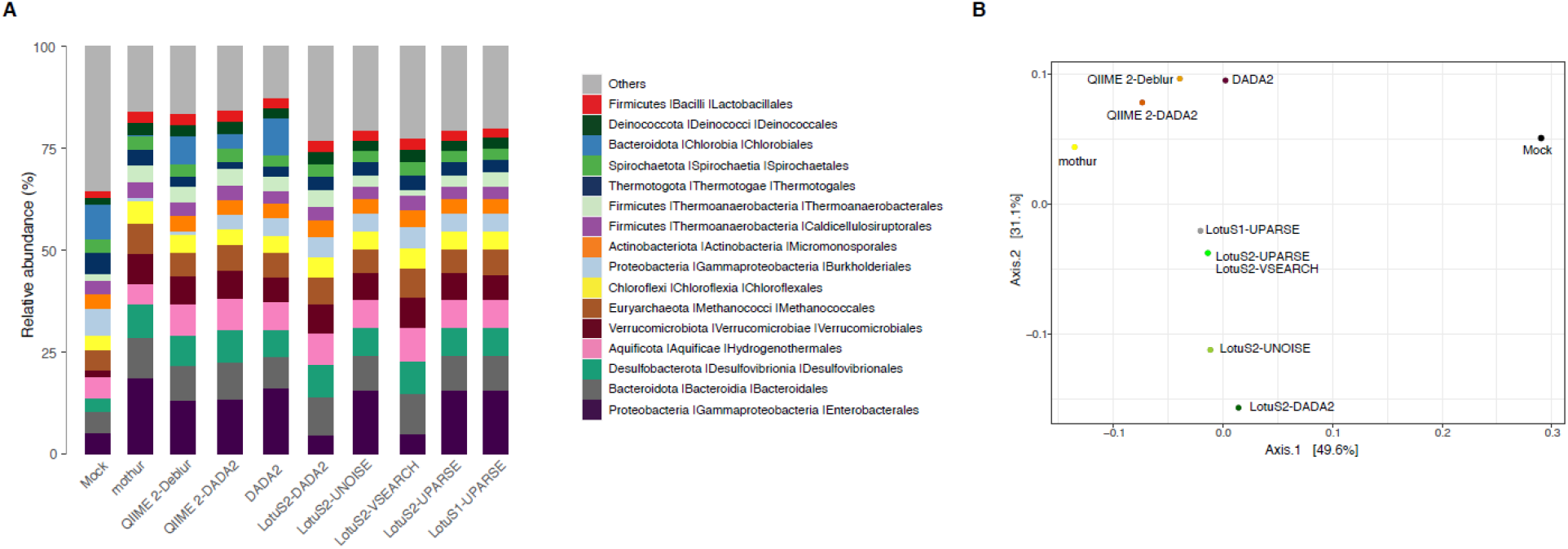
Observed composition of the mock community compared to the composition predicted by each pipeline. A) Relative abundances of the 16 orders having the highest abundance. B) Bray-Curtis distance based PCoA of the observed composition of the mock sample and composition predicted by each pipeline

**Supp. Figure 8:**
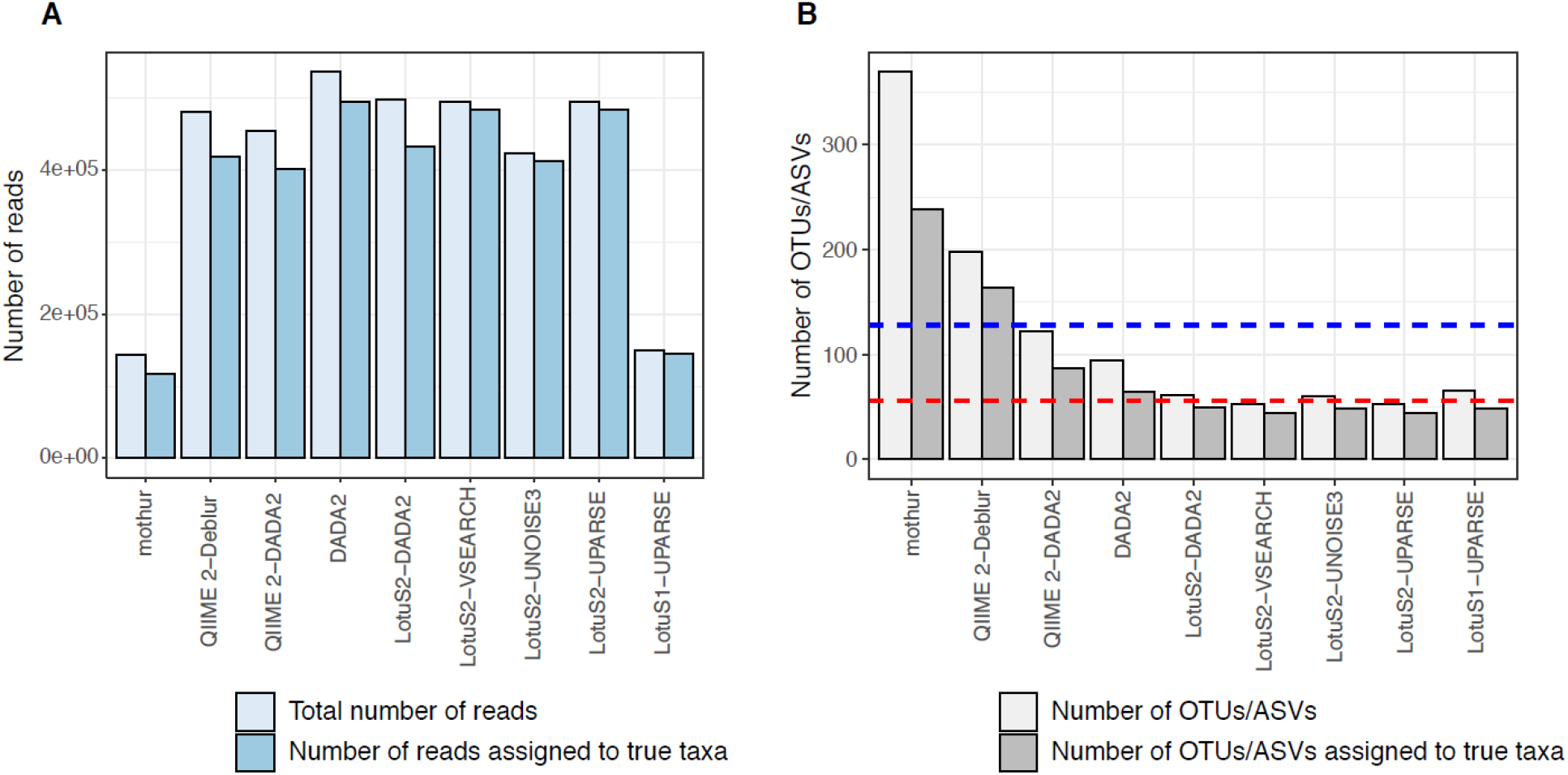
Number of reads and OTUs/ASVs and those assigned true taxa at genus level by each pipeline in the analysis of the mock community. Total number of A) reads retrieved by each pipeline and those assigned to true taxa at genus level B) OTUs/ASVs generated by each pipeline and those assigned to true taxa at genus level. Blue and red line indicates number of 16S gene copies and species, respectively, in the mock community.

**Supp. Figure 9:**
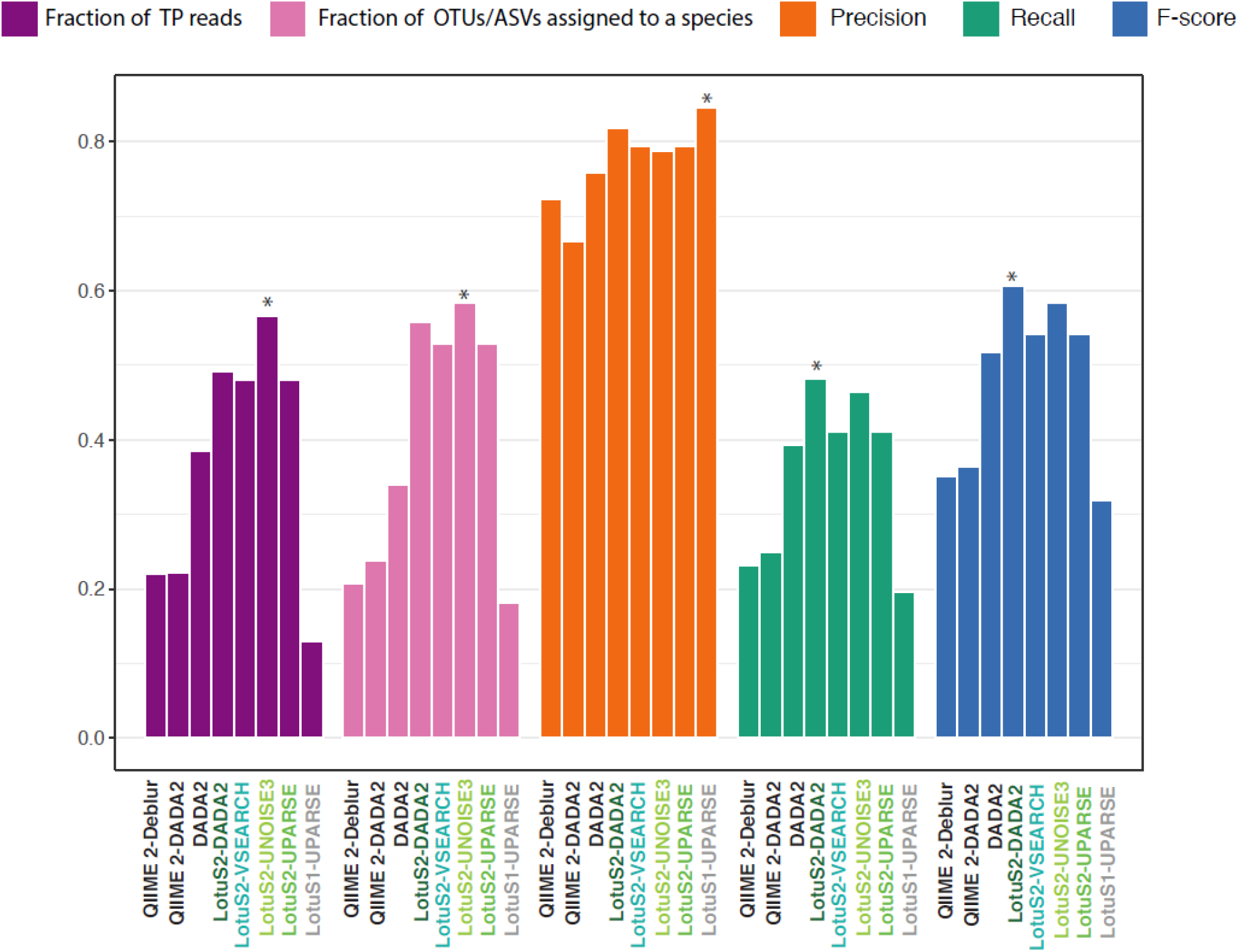
Accuracy of each pipeline in predicting the mock community composition at species level. For benchmarking we compared the fraction of reads assigned to true taxa and both correctly and erroneously recovered taxa at the species level from the mock community.

**Supp. Table 1:** Read counts and number of OTUs/ASVs in the OTU/ASV matrix of each pipeline.

| Gut-16S |  |  |
| --- | --- | --- |
|  | Number of reads | Number of OTUs/ASVs |
| mothur | 11855762 | 23736 |
| QIIME 2-Deblur | 9995254 | 950 |
| QIIME 2-DADA2 | 11510552 | 2539 |
| DADA2 | 12048048 | 2591 |
| LotuS2-DADA2 | 12935664 | 999 |
| LotuS2-UNOISE3 | 3698064 | 766 |
| LotuS2-UPARSE | 12995784 | 742 |
| LotuS2-VSEARCH | 2778696 | 1464 |
| LotuS1-UPARSE | 1305288 | 514 |

| Soil-16S |  |  |
| --- | --- | --- |
|  | Number of reads | Number of OTUs/ASVs |
| QIIME 2-Deblur | 1157357 | 19641 |
| QIIME 2-DADA2 | 2278731 | 25229 |
| DADA2 | 4526920 | 49111 |
| LotuS2-DADA2 | 2710629 | 19568 |
| LotuS2-UNOISE3 | 2448475 | 19217 |
| LotuS2-UPARSE | 2637572 | 8789 |
| LotuS2-VSEARCH | 2678716 | 9250 |
| LotuS1-UPARSE | 749449 | 5987 |

| Soil-ITS |  |  |
| --- | --- | --- |
|  | Number of reads | Number of OTUs/ASVs |
| QIIME 2-DADA2 | 4962260 | 2203 |
| DADA2 | 1742895 | 3368 |
| LotuS2-UPARSE | 4805387 | 1046 |
| LotuS2-VSEARCH | 4829288 | 920 |
| LotuS2-CDHIT | 2678716 | 1008 |

**Supp. Table 2: Significance of differences between each pipeline in the reproducibility of beta diversity between the technical replicates**

Significance of differences in Bray-Curtis distance between the pipelines were calculated based on the Tukey’s HSD test.

**Supp. Table 3:** Correlation and beta distance between the mock community and reconstructed mock community by each pipeline. **A-B)** Spearman and Pearson correlation between the expected abundances in the mock community and the observed abundances by each pipeline. **C)** Bray-Curtis dissimilarity between the known mock community and re-constructed mock community composition by each pipeline.

| Spearman Correlation |  |  |
| --- | --- | --- |
|  | p.value | correlation coefficient |
| mothur | 1.83E-07 | 0.544018417 |
| QIIME 2-Deblur | 1.57E-15 | 0.747912391 |
| QIIME 2-DADA2 | 3.76E-12 | 0.680648974 |
| DADA2 | 6.77E-12 | 0.674725632 |
| LotuS2-DADA2 | 3.26E-12 | 0.682064113 |
| LotuS2-VSEARCH | 2.80E-17 | 0.776030912 |
| LotuS2-UNOISE3 | 4.99E-14 | 0.720369663 |
| LotuS2-UPARSE | 2.80E-17 | 0.776030912 |
| LotuS2-UPARSE | 1.32E-19 | 0.808037907 |

| Pearson Correlation |  |  |
| --- | --- | --- |
|  | p.value | correlation coefficient |
| mothur | 3.99E-07 | 0.531185654 |
| QIIME 2-Deblur | 1.99E-11 | 0.663501229 |
| QIIME 2-DADA2 | 3.91E-09 | 0.600486282 |
| DADA2 | 7.72E-12 | 0.673389135 |
| LotuS2-DADA2 | 6.62E-05 | 0.43083946 |
| LotuS2-VSEARCH | 2.68E-09 | 0.605505625 |
| LotuS2-UNOISE3 | 1.22E-08 | 0.584843731 |
| LotuS2-UPARSE | 2.68E-09 | 0.605505625 |
| LotuS1-UPARSE | 1.63E-09 | 0.611973422 |

| BCd to the mock community |  |
| --- | --- |
|  | BCd |
| mothur | 0.430087 |
| QIIME 2-Deblur | 0.340823 |
| QIIME 2-DADA2 | 0.373356 |
| DADA2 | 0.327616 |
| LotuS2-DADA2 | 0.35983 |
| LotuS2-VSEARCH | 0.324378 |
| LotuS2-UNOISE3 | 0.34578 |
| LotuS2-UPARSE | 0.324378 |
| LotuS1-UPARSE | 0.324448 |

**Supp. Table 4:** Accuracy of each pipeline in re-constructing the mock community at genus level.

|  | Number of OTUs/ASVs | Number of reads | Fraction of reads assigned to TP taxa | TP | FP | FN | Precision | Recall | F-score |
| --- | --- | --- | --- | --- | --- | --- | --- | --- | --- |
| mothur | 370 | 144147 | 0.817443304 | 34 | 25 | 11 | 0.576271 | 0.755556 | 0.653846 |
| QIIME 2-Deblur | 198 | 480049 | 0.872517181 | 38 | 13 | 7 | 0.745098 | 0.844444 | 0.791667 |
| QIIME 2-DADA2 | 122 | 454082 | 0.882792095 | 36 | 19 | 9 | 0.654545 | 0.8 | 0.72 |
| DADA2 | 94 | 536901 | 0.922646819 | 39 | 24 | 6 | 0.619048 | 0.866667 | 0.722222 |
| LotuS2-DADA2 | 61 | 497970 | 0.867775167 | 38 | 9 | 7 | 0.808511 | 0.844444 | 0.826087 |
| LotuS2-VSEARCH | 53 | 494122 | 0.979268278 | 39 | 9 | 6 | 0.8125 | 0.866667 | 0.83871 |
| LotuS2-UNOISE3 | 60 | 423292 | 0.975794487 | 38 | 9 | 7 | 0.808511 | 0.844444 | 0.826087 |
| LotuS2-UPARSE | 53 | 494122 | 0.979268278 | 39 | 9 | 6 | 0.8125 | 0.866667 | 0.83871 |
| LotuS1-UPARSE | 66 | 148959 | 0.979202331 | 41 | 16 | 4 | 0.719298 | 0.911111 | 0.803922 |

## Supplementary Information

### Influence of dereplication thresholds, non-default parameters and read truncation

Dereplication is the pre-clustering of sequencing reads at 100% nucleotide identity, a commonly used strategy to reduce the computational complexity of sequence clustering [17]. Further, dereplication can be used to filter out sparsely occurring reads that could represent technical artifacts, unlikely to represent true biodiversity. Therefore, LotuS2 uses a “dereplication” filter, that can be user defined.

Overall, this filter does not mostly change the number of OTU/ASV counts, with more OTUs/ASVs being recovered when the filter is more relaxed **(Supp. Figure 2A,D,G)**. This is expected because this filter is designed to remove sparse OTUs/ASVs that could both represent technical replicates as well as extremely rare microbes. However, this did not affect the overall community reproducibility of either gut- or soil-16S samples. However, in soil-ITS samples, we noted a dramatic decrease in BCd between technical replicates at stricter dereplication cut-offs **(Supp. Figure 2H-I)**.

The number of retrieved reads remained very stable independent of filtering stringency; this is expected because the backmapping of mid-quality reads will re-introduce reads not passing the dereplication filter.

LotuS2 uses several default options (-lulu 1, -xtalk 0, -keepUnclassified 0 and -ITSX 1; where “1” means the option is “activated” and “0” means “deactivated”). When activated, -lulu option uses LULU R package [23] to merge OTUs/ASVs based on their co-occurrences; -xtalk option checks for cross-talk [32], -keepUnclassified includes unclassified (i.e. not matching to any taxon in the taxonomy database) OTUs/ASVs in the final matrix and –ITSx activates the ITSx program [31] to only retain OTUs fitting to ITS1/ITS2 hmm models. The impact of these parameters on the reproducibility of LotuS2 was tested **(Supp. Figure 3)**. Overall, non-default options did not change the BCd between the technical replicates except -keepUnclassified 1 notably increasing BCd in gut-16S, while -lulu 0 slightly increased BCd in soil-ITS.

Read length truncation is frequently used to remove the typically low quality 3’ end of reads [8,17]. This is impacting the retrieved read counts as well as observed OTU/ASV diversity. For example, at 170 bp read truncation, mothur, DADA2 and QIIME 2-DADA2 were severely impacted in merging read pairs, failing or only integrating a fraction of read pairs in gut and soil-16S datasets **Supp. Figure 4**). While LotuS2 also had slightly different read and cluster numbers with changing truncation lengths, it was more stable, because reads are merged in the seed extension step after sequence clustering on truncated, high-quality reads are completed **(Supp. Figure 4)**. In shorter or longer read truncations, LotuS2 was still performing the best with the lowest BCd **(Supp. Figure 5A,C)** and Jd **(Supp. Figure 5B,D)** between technical replicates in both gut- and soil-16S datasets.

Taken together, the higher performance of LotuS2 in reproducibility of the dataset was independent of the dereplication parameters and read truncation length.

## Footnotes

1 Note that UNOISE3 uses the term zero-range OTUs (zOTUs); for brevity, this is omitted throughout the text.

